# A Pro-Diabetogenic mtDNA Polymorphism in the Mitochondrial-Derived Peptide, MOTS-c

**DOI:** 10.1101/695585

**Authors:** Hirofumi Zempo, Su-Jeong Kim, Noriyuki Fuku, Yuichiro Nishida, Yasuki Higaki, Junxiang Wan, Kelvin Yen, Brendan Miller, Roberto Vicinanza, Eri Miyamoto-Mikami, Hiroshi Kumagai, Hisashi Naito, Jialin Xiao, Hemal H. Mehta, Changhan Lee, Megumi Hara, Yesha M. Patel, Veronica W. Setiawan, Timothy M. Moore, Andrea L. Hevener, Yoichi Sutoh, Atsushi Shimizu, Kaname Kojima, Kengo Kinoshita, Keitaro Tanaka, Pinchas Cohen

**Author notes:** Co-first authors (Hirofumi Zempo, Su-Jeong Kim). Share senior authorship (Keitaro Tanaka, Pinchas Cohen). Correspondence to Noriyuki Fuku, PhD, Pinchas Cohen, MD.

## Abstract

Type 2 Diabetes (T2D) is an emerging public health problem in Asia. An Asian mitochondrial DNA variation m.1382A>C (rs111033358) leads to a K14Q amino acid replacement in MOTS-c, an insulin sensitizing mitochondrial-derived peptide. Meta-analysis of three cohorts (n=27,527, J-MICC, MEC, and TMM) showed that males but not females with the C-allele exhibit a higher prevalence of T2D. Furthermore, in J-MICC, only males with the C-allele in the lowest tertile of physical activity increased their prevalence of T2D, demonstrating a kinesio-genomic interaction. High-fat fed, male mice injected with MOTS-c showed reduced weight and improved glucose tolerance, but not K14Q-MOTS-c treated mice. Like the human data, female mice were unaffected. Mechanistically, K14Q-MOTS-c leads to diminished insulin-sensitization *in vitro*. Thus, the m.1382A>C polymorphism is associated with susceptibility to T2D in men, possibly interacting with exercise, and contributing to the risk of T2D in sedentary males by reducing the activity of MOTS-c.

## INTRODUCTION

The prevalence of type 2 diabetes mellitus (T2D) is growing dramatically. Over 400 million individuals were diagnosed with T2D worldwide in 2015, and the International Diabetes Federation projects that at least 600 million people will need to be treated for T2D by 2040 ^1^. However, treating T2D is challenging because the disease etiology is genetically heterogeneous and varies among ethnicities ^2–4^. For example, a recent meta-analysis showed that T2D is 72% heritable (95% CI: 61-78%) ^2^, and the Western Pacific Region – including China and Japan – makes up ∼37% of the total T2D diagnoses in the world ^1^. This is particularly noteworthy because although East-Asian populations have lower mean body mass index (BMI) than Caucasian populations, they have a higher susceptibility to T2D compared to Caucasians ^5^. Furthermore, when matched for BMI, East Asians have a greater percentage of body fat and a tendency for increased visceral adiposity ^6^. T2D in Asian patients is characterized by early beta cell dysfunction, in contrast to Caucasian populations that have more insulin resistance ^7, 8^. Thus, ethnicity-based DNA variations likely influence the pathogenesis of T2D.

While diabetes directly caused by mutations in mtDNA are extremely rare ^9^, several genetic analyses revealed that mtDNA polymorphisms contribute to T2D risk in both European and Asian populations ^10, 11^. Notably, mtDNA sequences are more varied by ethnicity compared to nuclear DNA sequences due to the 10-times higher mutation rate of mtDNA compared to nuclear DNA ^12, 13^. As a result, certain mtDNA polymorphisms could contribute to the T2D prevalence in different ethnicities. The human mitochondrial genome consists of 16,569-base pairs and encodes 37 genes including 13 full size proteins involved in oxidative phosphorylation, 2 rRNA, and 22 tRNA genes that are necessary for protein synthesis within the mitochondria. Genetic polymorphisms in the mtDNA could potentially affect cell metabolism ^14^ leading to alterations in insulin signaling or beta cell function and may explain why T2D prevalence is different between ethnicities.

Recent studies showed that in addition to the above mentioned 37 genes, the mitochondrial chromosome also encodes multiple biologically active peptides derived from small open reading frames (ORF) ^15–19^. We discovered one such mitochondrial derived peptide (MDP) encoded from the 12S rRNA in the mtDNA called <u>m</u>itochondrial <u>o</u>pen-reading-frame of the twelve <u>S</u> rRNA –<u>c</u> (MOTS-c) ^20^. MOTS-c is a 16-amino acid peptide expressed in multiple tissues, including skeletal muscles, and is detected in the plasma. MOTS-c activates AMPK and acts as an insulin sensitizer ^18, 21, 22^. MOTS-c regulates insulin sensitivity and metabolic homeostasis in the skeletal muscle of mice ^22^. MOTS-c reduces weight gain and decreases fat accumulation in the liver in high-fat diet-induced obese mice, leading some to calling it an “exercise-mimetic peptide” ^22, 23^. MOTS-c prevents ovariectomy-induced metabolic dysfunction ^24, 25^. MOTS-c levels are correlated with insulin resistance, and circulating MOTS-c levels are reduced in obese male (but not female) children ^26, 27^. MOTS-c levels are inversely correlated with markers of insulin resistance and obesity including BMI, waist circumference, waist-to-hip ratio, fasting insulin level, HOMA-IR, HbA_1c_ ^27^. In addition, MOTS-c levels are positively correlated with endothelial function in humans ^28^ and MOTS-c administration confers protection from ovariectomy-induced osteoporosis in mice via the AMPK pathway ^21^.

An East Asian-specific mtDNA Single Nucleotide Polymorphism (SNP), m.1382A>C (rs111033358), causes an amino acid replacement from Lys (K) to Gln (Q) at the 14th amino-acid residue in the MOTS-c peptide ^29^. This SNP was noted in Japanese with exceptional lifespan ^29^. However, the relation between the m.1382A>C polymorphism and human T2D pathophysiology remains unknown. The purpose of this study was two-fold: 1) elucidate the association between T2D and m.1382A>C in Japanese individuals, and 2) establish the biological effects of the K14Q-MOTS-c peptide variant – a consequence of the m.1382A>C polymorphism – on insulin action and adiposity *in vitro* and *in vivo*.

## RESULTS

### A MOTS-c SNP affects T2D incidence in Japanese men

We examined the effect of the m.1382A>C polymorphism (rs111033358), which is imbedded within the ORF of the MOTS-c peptide on T2D in three cohorts of individuals of Japanese descent. First, we used the Japan Multi-Institutional Collaborative Cohort Study (J-MICC), which includes 4,963 men and 6,889 women, of whom 98.2% had documented BMI, smoking status, diagnosis of T2D, m.1382A>C polymorphism status (Supplemental Table 1). Second, we used Japanese-American subjects in the Multiethnic Cohort (MEC) study, which includes 1,810 men and 1,577 women of Japanese descent living in the US. Third, we used the Tohoku Medical Megabank project (TMM), which includes 4,471 males and 7,817 females (Supplemental Table 2). Meta-analysis of the three cohorts showed that the m.1382A>C polymorphism significantly increases the prevalence of T2D (*p* < 0.01) in males but not in females (Figure 1A and 1B). In fact, in the MEC study men with the C allele showed increase in T2D risk (*p* = 0.04) compared to A allele carriers (Supplemental Table 3).

**Figure 1.**
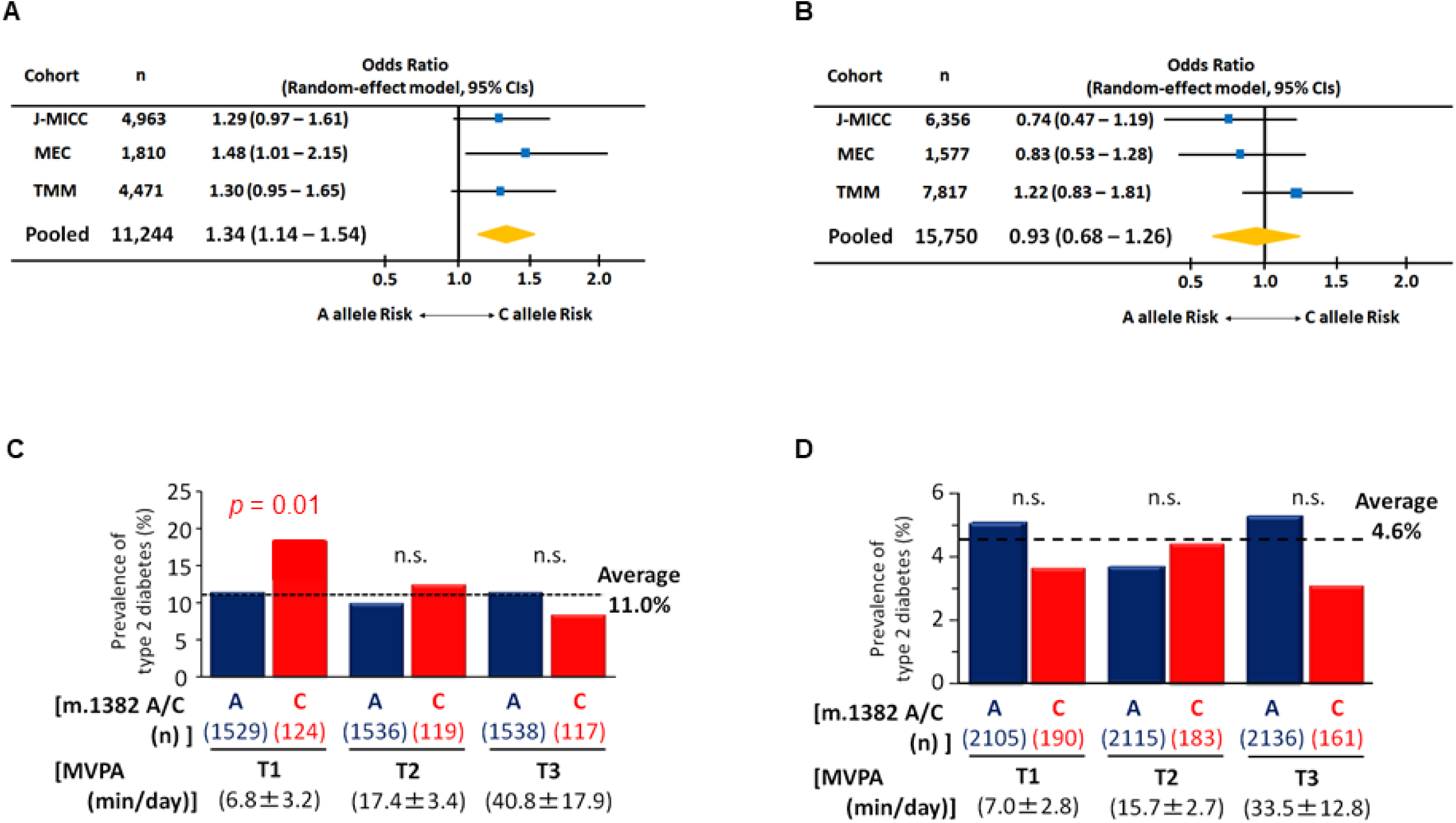
The prevalence of type 2 diabetes in m.1382A>C polymorphism. (A) Forest plot for meta-analysis (pooled random effects) on the prevalence of type 2 diabetes in males. Odds ratios are adjusted by age and BMI. Test for the pooled effect, Z = 2.86, *p* < 0.01. Heterogeneity was insignificant (I^2^ = 0%, *p* = 0.84). (B) Forest plot for meta-analysis (pooled random effects) on the prevalence of type 2 diabetes in female. Odds ratios are adjusted by age and BMI. Test for the pooled effect, Z = −0.46, *p* = 0.64. Heterogeneity was insignificant (I^2^ = 33.6%, *p* = 0.22). In Japan Multi-Institutional Collaborative Cohort Study (J-MICC) study, m.1382A>C polymorphism carriers divided by tertile of physical activity in (C) males and (D) females. Activity was assessed by the degree of moderate-to-vigorous intensity physical activity (MVPA). In men only, sedentary levels of MVPA were associated with an increased risk of diabetes in the C allele. Cochran-Armitage trend test of m.1382 C allele for men, χ^2^ = 6.26, *p* = 0.012. Cochran-Armitage trend test of m.1382 A allele for females, χ^2^ = 0.03, *p* = 0.861.

### A Kinesio-Genomic effect of a MOTS-c SNP on T2D incidence in men

The m.1382A>C polymorphism increases T2D risk in sedentary males in J-MICC. Men with the C allele of the m.1382A>C polymorphism showed a trend for higher risk for T2D regardless of physical activity status measured by accelerometer (C allele, 13.1%; A allele, 10.8%; *p* = 0.196, Supplemental Table 1 top). However, when assessing potential contributors to T2D risk, the degree of physical activity, measured by accelerometer, and expressed as moderate-to vigorous-intensity physical activity (MVPA), correlated with lower risk for T2D in men with the C allele (MVPA beta = 0.023; *p* = 0.035; Tables 1-2). As shown in Figure 1C, men with the C allele and low physical activity (T1 of MVPA: 6.8 ± 3.2 min/day) had a 65% greater rate of T2D than men with the A allele and low physical activity (A allele T2D prevalence = 11.2%; C allele T2D prevalence = 18.5%; *p* = 0.014; Figure 1C). This relationship is not seen in men with higher physical activity. The effect of m.1382A>C is male-specific, as females with the C allele who were classified as sedentary (physical activity T1 = 7.0 ± 2.8 min/day) did not have an increased T2D risk (A allele T2D prevalence = 5.1%; C allele T2D prevalence = 3.7%; *p* = 0.258; Figure 1D). No significant differences were found for other characteristics between the m.1382A>C genotype in men with the lowest MVPA (Supplemental Table 4). These results strongly suggest that a combination of sedentary lifestyle and the m.1382A>C polymorphism contributes to elevated T2D risk. Unfortunately, the TMM study and MEC study only quantified physical activity using a questionnaire, which is different from the accelerometer-based measurement used by J-MICC. Physical activity questionnaires do not correlate well with accelerometer-based measurements in young and old individuals ^30–33^. Hence, we are unable to assess physical activity behavior among the three cohorts using a harmonized approach (i.e., accelerometer). As a consequence, it appears that the kinesio-genomic relationship of m.1382A>C genotype and T2D can only be teased out with accurate exercise assessment in the J-MICC cohort.

**Table 1.**
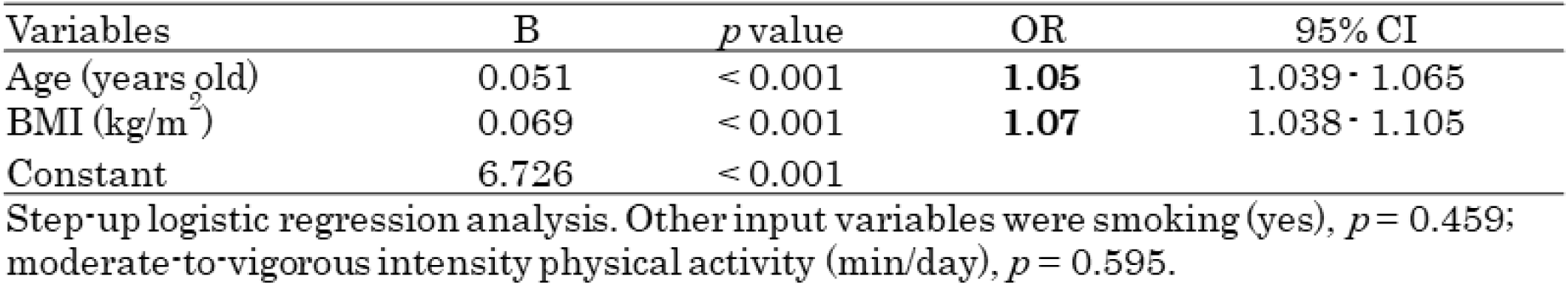
Factors associated with the risk of type 2 diabetes in male with the m.1382 A allele in J-MICC.

**Table 2.**
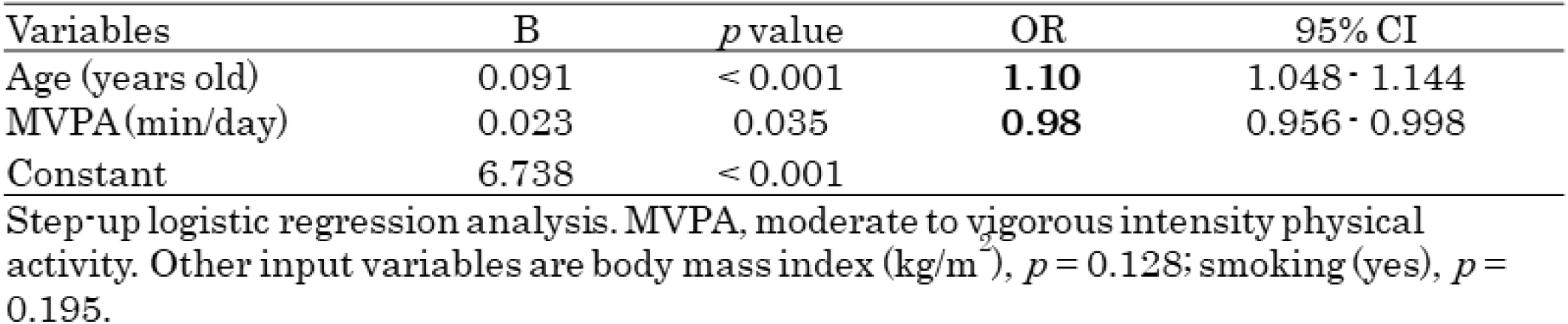
Factors associated with the risk of type 2 diabetes in male with the m.1382 C allele in J-MICC.

### Effects of other polymorphisms on the incidence of T2D

We assessed the effect of m.1382A>C on T2D independent from separate polymorphism-determining haplogroups D, D4 and D4b by using J-MICC cohort. The m.1382A>C polymorphism is one of the determinant polymorphisms in the mitochondrial haplogroup D4b2. This haplogroup is most frequent among East Asians including Japanese, Koreans, and Northern Chinese. Korean men in haplogroup D4b have previously been shown to have a dramatically increased risk of T2D (OR 3.55 [95% CI 1.65–8.34], *p* = 0.0019) ^11^. We analyzed four additional SNPs defining haplogroup D: m.4883C>T (ND2: syn), m.5178C>A (ND2: L237M), haplogroup D4: m.3010G>A (16S rRNA), and haplogroup D4b: m.15440T>C (CYB: D4b1) and m.1382A>C (D4b2) for an effect on T2D risk (Supplemental Tables 5-8). None of these SNPs were associated with T2D risk; only m.1382A>C significantly increased T2D prevalence in sedentary male subjects (Supplemental Table 9). We did not look at the effect of m. 8020G>A (D4b) because 1) m. 8020G>A status was not collected in J-MICC and 2) The polymorphism m.8020G>A is not only a SNP of haplogroup D4b but also a SNP of other haplogroups (i.e., M7b1a1a1a, D6a1a, R0a1a2, H5k, J1c1g1, F2e1, F4b and K1a14). The D4b haplogroup (defined as possessing both the m.15440 and m.1382 polymorphisms) and the D4b1 determinant SNP, m.15440, also does not show any effect on the risk of T2D. A previous study reported that haplogroup D4b was associated with T2D in Korean men, but not women. We hypothesized that the increase of risk in Korean male carriers of haplogroup D4b is driven by the m.1382A>C polymorphism. Because m.1382A>C defining sub-haplogroup D4b2 is a major component of haplogroup D4b, it is likely that this polymorphism influences the increased risk in T2D in Korean men due to the resultant MOTS-c K14Q variant. We speculate that the increased risk in Korean male carriers of haplogroup D4b is driven by the m.1382A>C polymorphism because D4b haplogroup itself does not impact T2D risk in J-MICC study.

### 3D structure of WT and K14Q-MOTS-c

The m.1382A>C polymorphism is located in the 12S rRNA within the mtDNA, which encodes the ORF for the MOTS-c peptide. Therefore, the m.1382A>C polymorphism could alter both the 12S rRNA structure and causes a K14Q replacement in the MOTS-c peptide. We modeled the 12S rRNA structure by m.1382A and m.1382C and found very little difference between the two models (Figure 2A). Next, we examined the difference between WT-MOTS-c (K14) and K14Q-MOTS-c (Q14). The electro physical differences predict 14^th^ residue of WT-MOTS-c is positively charged, whereas same position of K14Q-MOTS-c is neutral (Figure 2B). The mPROVEAN (PROtein Variation Effect Analyzer; http://provean.jcvi.org) analyses revealed that the polymorphism could be biologically relevant; that is, the m.1382A>C polymorphism on MOTS-c that makes K14Q-MOTS-c had a score of −4.0, below the specifically predicted cutoff score (= −2.5), above which the variant would be “neutral” ^34^. We compared the structure of WT and K14Q-MOTS-c. The 3D structures of WT- and K14Q-MOTS-c were predicted via the de novo modeling servers, PEP-FOLD3 and I-TASSER (Model 1, Supplemental Figure 1A and 1B; Model 2, Supplemental Figure 1C and 1D). The analysis indicated that the 3D structure of K14Q-MOTS-C is substantially different compared to WT-MOTS-c.

**Figure 2.**
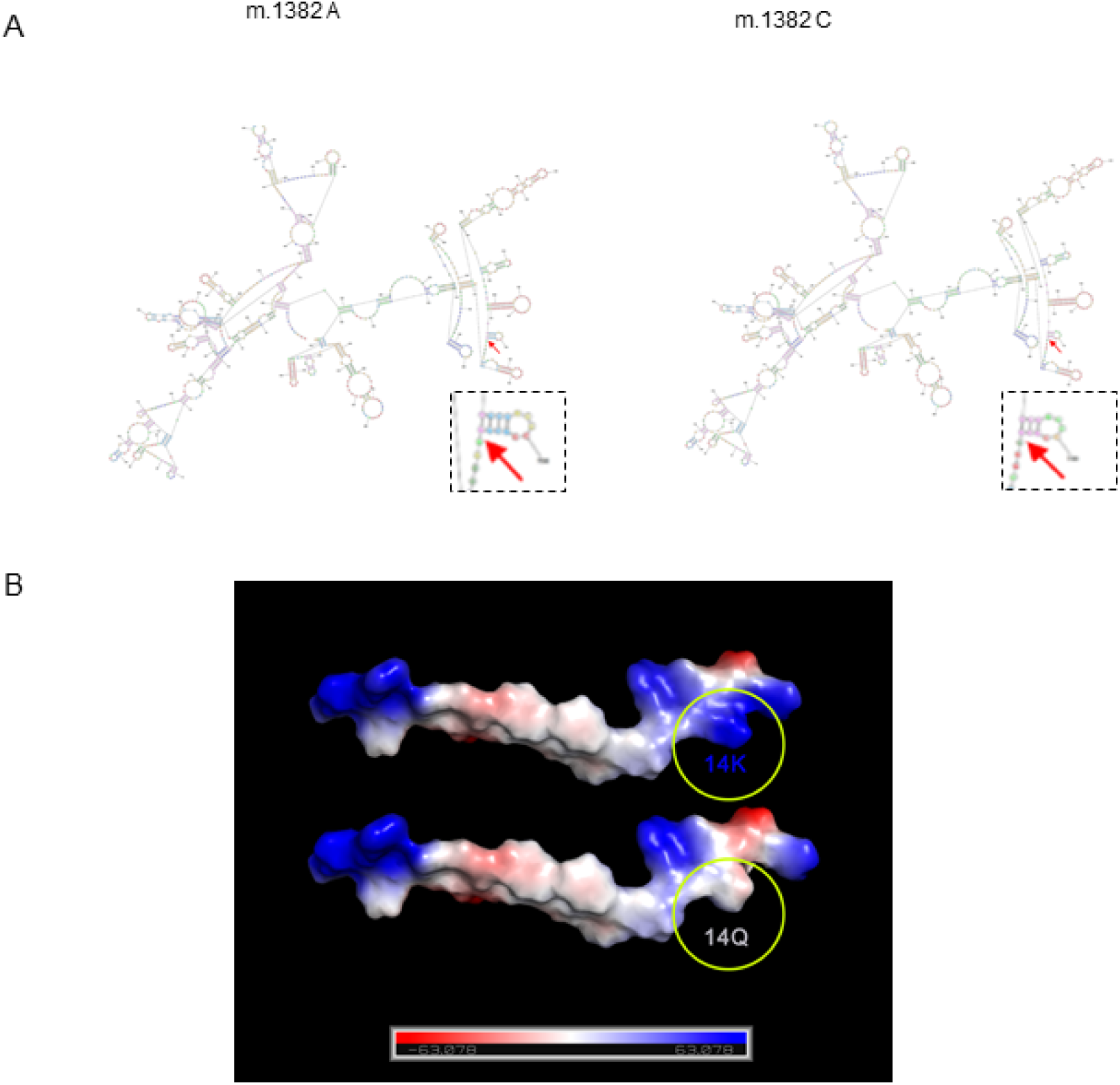
The impact of m.1382 A>C polymorphism in the structure of 12S rRNA and MOTS-c peptide. (A) The predicted 12S rRNA structure by m.1382 A>C polymorphism by RNAstructure (https://rna.urmc.rochester.edu/RNAstructureWeb/Servers/Predict1/Predict1.html). The box is the enlarged view of the pointed with arrow. (B) The electro physical differences between m.1382 A and C analyzed by PyMOL.

As MOTS-c has beneficial effects on metabolism and acts as an exercise mimetic, we speculated that sedentary men who carry the C allele, which produces the K14Q-MOTS-c peptide will have a reduced insulin-sensitizing effect compared to WT-MOTS-c in cellular and animal models.

### Insulin sensitizing effects of WT- and K14Q-MOTS-c *in vitro*

Since MOTS-c has been shown to increase insulin sensitivity in L6 myotubes and mouse skeletal muscle via glucose clamps and insulin action assays, we assessed the effect of K14Q-MOTS-c on cellular insulin action by exogenous peptide administration and by transient transfection. As exogenous MOTS-c treatment in mice showed a substantial effect to improve insulin action and since peptide treatments are clinically translatable, we treated cells with exogenous WT- and K14Q-MOTS-c. To examine the insulin sensitizing effect of MOTS-c and its variant, we treated cells with insulin in the absence or presence of WT- and K14Q-MOTS-c in differentiated C2C12 myotubes. Upon insulin stimulation, insulin receptor tyrosine kinase activates AKT via PI3K. AKT phosphorylation by insulin is higher in WT-MOTS-c than control and K14Q-MOTS-c treatment in differentiated C2C12 myotubes (Figure 3A). WT-MOTS-c further increased AKT phosphorylation in the presence of insulin, whereas K14Q-MOTS-c showed a blunted response to insulin-stimulated AKT phosphorylation in the presence of insulin in differentiated C2C12 myotubes (Figure 3A). In addition, insulin plays an important role in adipocyte differentiation. Hence, we used 3T3-L1 pre-adipocyte differentiation as another model to assess insulin sensitivity. During 3T3-L1 pre-adipocyte differentiation, cells were incubated with insulin (1 µg/mL) containing medium. We treated cells with insulin (0, 0.5, and 1 µg/mL) in the presence or absence of WT-MOTS-c. MOTS-c increased lipid droplets only in the presence of insulin, suggesting MOTS-c has an insulin sensitizing effect on adipocyte differentiation (*data not shown*). Next, we treated cells with WT and K14Q-MOTS-c in the presence of 1µg/mL insulin. WT-MOTS-c, but not K14Q-MOTS-c increased lipid droplets during adipocyte differentiation (Figure 3B). Next, differentiated L6 myotubes and 3T3-L1 cells were treated with WT and K14Q-MOTS-c. Glucose levels were measured in the conditioned medium. WT-MOTS-c increased glucose uptake, whereas K14Q-MOTS-c was less effective at enhancing insulin-stimulated glucose uptake than WT-MOTS-c (Figure 3C and D). We also transiently transfected HEK293 cells with WT-MOTS-c and K14Q-MOTS-c and measured glucose uptake. WT-MOTS-c increased glucose uptake, whereas K14Q-MOTS-c effects were the same as the control plasmid (Figure 3E). Taken together, our *in vitro* studies show that K14Q-MOTS-c has a diminished action as an insulin sensitizer compared to WT-MOTS-c.

**Figure 3.**
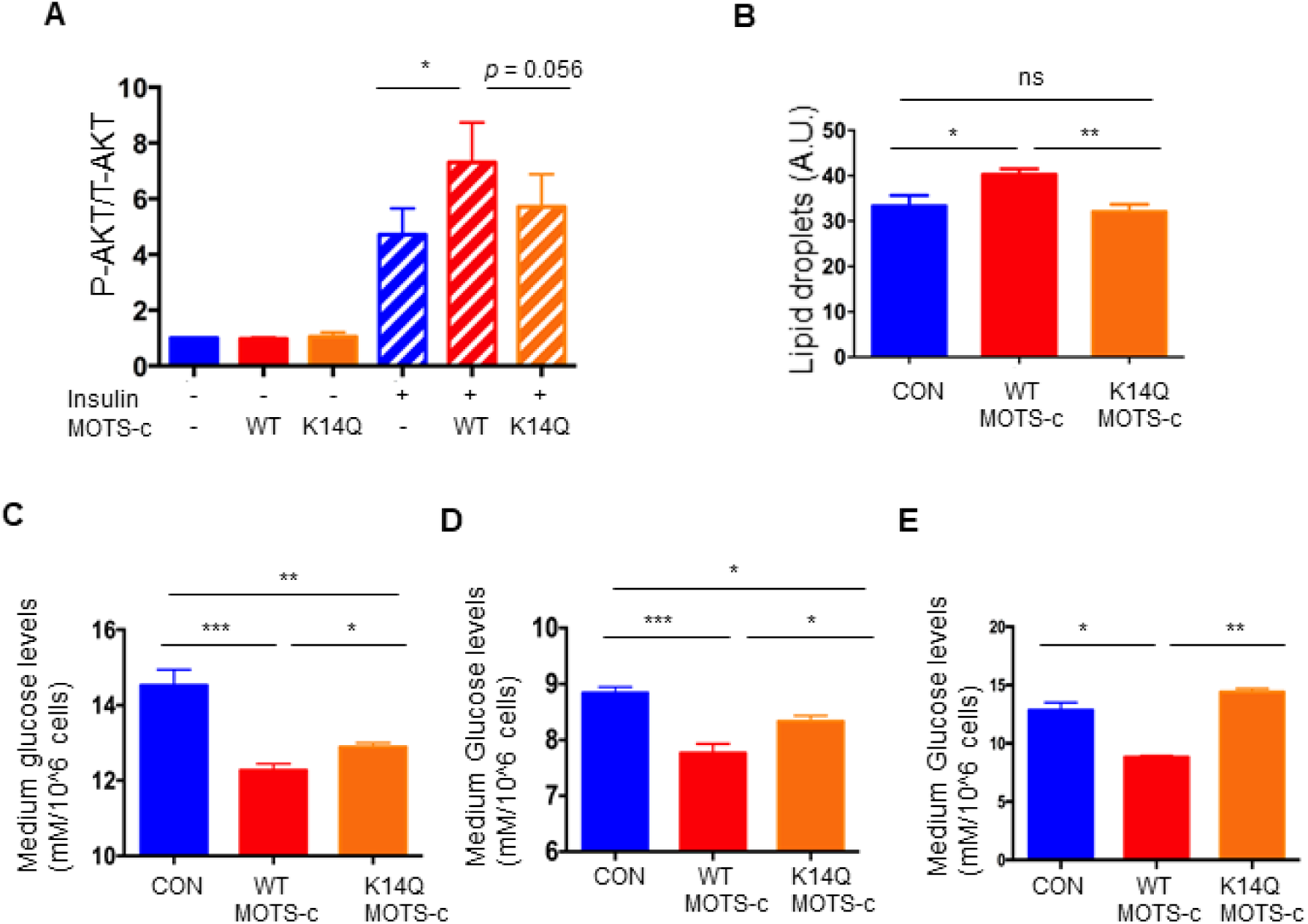
K14Q MOTS-c has a reduced insulin sensitizing effect compared to WT MOTS-c in muscle and fat. (A) Differentiated C2C12 myotubes, were treated with WT and K14Q MOTS-c in the absence or presence of 10-nM insulin. The total and phosphorylated AKT (Ser473) levels were measured by MSD. (B) 3T3L1 Cells were incubated with 1-µg/mL insulin +/- WT MOTS-c or K14Q MOTS-c. At day 12, lipid droplets were measured by Nile Red staining. (C) Differentiated L6 myotubes and (D) differentiated 3T3-L1 were treated with WT and K14Q MOTS-c for 48hr. Glucose levels in the medium were measured. (E) Overexpression of WT MOTS-c, but not K14Q, increases glucose uptake in HEK293 Human embryonic kidney cells. HEK293 cells were transiently transfected with WT MOTS-c and K14Q MOTS-c for 72 hr. Data are reported as mean ± SEM of seven independent experiments. *** *p*<0.001** *p*<0.01, \**p*<0.05.

### WT-MOTS-c and K14Q-MOTS-c have differential effects on male mice fed a high fat diet

We administered WT and K14Q-MOTS-c (7.5-mg/kg; twice daily (BID); IP) to CD1 male mice fed a high-fat diet (HFD, 60% by calories) to examine the roles of K14Q-MOTS-c in diet-induced metabolic alterations. Our previous studies showed that MOTS-c reduced body weight gain and fat mass, improved insulin sensitivity in skeletal muscle, and heightened energy expenditure in HFD-fed CD1 mice. In this study, we confirmed our previous findings showing that WT-MOTS-c significantly reduced weight gain in high fat diet fed mice (Figure 4A). In contrast to WT-MOTS-c, K14Q-MOTS-c failed to protect against HFD-induced weight gain; thus, total body weight was identical between K14Q-MOTS-c and water-treated controls. (Figure 4A). Moreover, HFD feeding induced a similar fat weight gain between K14Q-MOTS-c and water-treated control, whereas WT-MOTS-c showed a reduced fat weight compared to control following the 3 weeks of HFD consumption (Figure 4B). There was no effect on lean mass in the HFD-fed mice treated with both WT and K14Q-MOTS-c (Figure 4C). Although body weight was lower in WT-MOTS-c treated group, food intake was identical between the three groups (Figure 4D). Since MOTS-c increased cellular glucose uptake and increased insulin sensitivity *in vitro,* we hypothesized its actions *in vivo* would be related to glucose clearance indicative of improved insulin sensitivity. We administrated mice with WT-MOTS-c and K14Q-MOTS-c for 21 days and then subjected them to a glucose tolerance test (GTT). We observed that WT-MOTS-c significantly enhanced glucose clearance, whereas the glucose tolerance curves was not different between K14Q-MOTS-c and water-treated group (Figure 4E and 4F). Taken together, K14Q-MOTS-c is less effective as a metabolic regulator in male mice *in vivo* compared to WT-MOTS-c.

**Figure 4.**
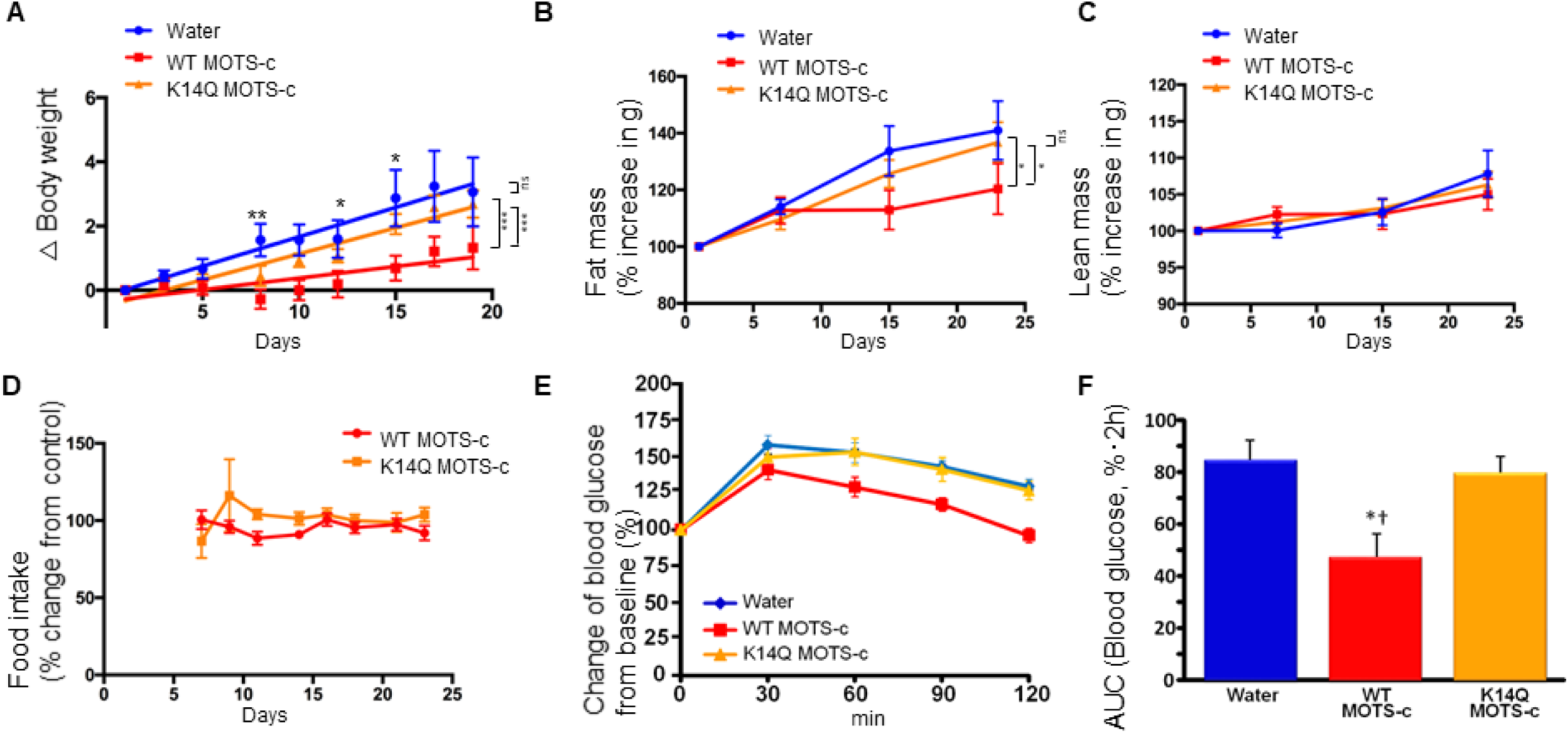
K14Q MOTS-c is less effective than WT MOTS-c, on reducing body weight, fat mass, glucose tolerance in CD1male mice exposed to high-fat diet. (A–D) Male CD-1 mice (14 weeks old) fed a high-fat diet (HFD, 60% by calories) (n = 8-12) treated with WT and K14Q MOTS-c (7.5 mg/kg; IP; BID) for 16 days. (A) body weight, (B) fat mass, (C) lean mass, (D) food intake. (E-F) Eight-weeks old male CD-1 mice fed a HFD diet (n =8-10) treated with WT and K14Q MOTS-c (0.5mg/kg; IP; daily) for 21 days. Then, they were tested glucose tolerance test (GTT). (E) Blood glucose and (F) glucose AUC, during in GTT. (A-D) *** *p*<0.001, ** *p*<0.01, \**p*<0.05. (E-F) \**p*<0.01 Water vs. WT MOTS-c. †*p*<0.01, K14Q MOTS- vs. WT MOTS-c.

### WT-MOTS-c and K14Q-MOTS-c have similar effects on female mice fed a high fat diet

We showed that the m.1382A>C polymorphism is associated with T2D in men but not in women. To identify whether this phenomenon can be recapitulated in mice, we administrated WT and K14Q-MOTS-c (7.5-mg/kg; BID; IP) in CD1 female mice fed a HFD (60% by calories). Unlike male mice, female mice did not respond to WT-MOTS-c treatment (Figure 5A). Similar to male mice, female mice injected with K14Q-MOTS-c also gained weight (Figure 5A). Fat mass, lean mass, and food intake were also not altered in WT and K14Q-MOTS-c injected female mice (Figure 5B-D). We administrated mice with WT-MOTS-c and K14Q-MOTS-c for 21 days and then subjected them to a glucose tolerance test (GTT). We observed that glucose curves for both WT-MOTS-c and K14Q-MOTS-c were not different from control (Figure 5E-F).

**Figure 5.**
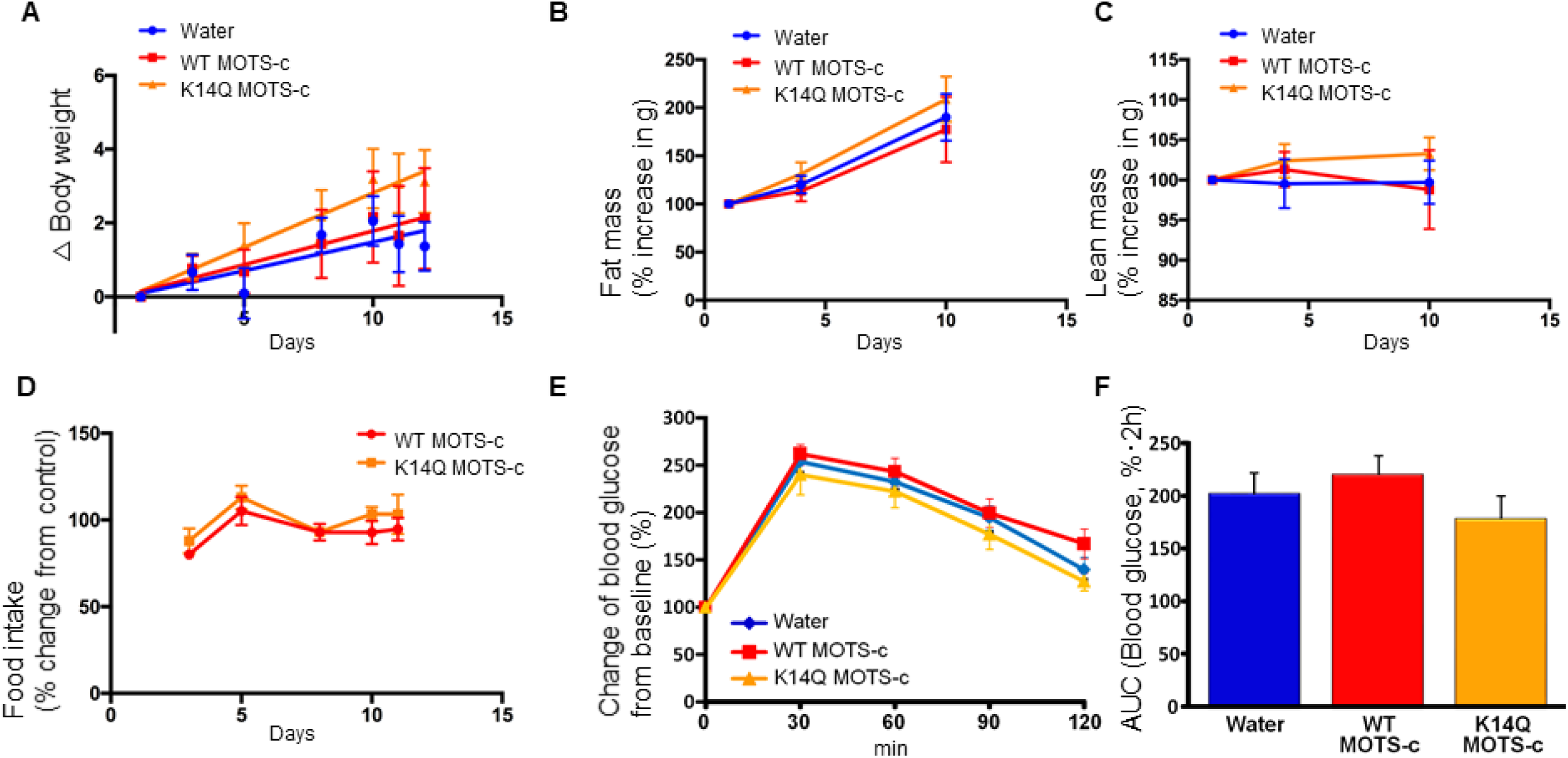
Diminished effects of MOTS-c in female mice. 12 weeks old CD1 mice fed a high fat diet were treated with WT MOTS-c and K14Q MOTS-c similarly to Figure 4A-D (n = 8) for 12 days. (A) body weight, (B) fat mass, (C) lean mass, (D) food intake. Eight-weeks old female mice fed a HFD diet (n =8-10) treated with WT and K14Q MOTS-c (0.5mg/kg; IP; daily) for 21 days. Then, they were tested glucose tolerance test (GTT). (E) Blood glucose and (F) glucose AUC, during in GTT.

### WT-MOTS-c and K14Q-MOTS-c have no effect on voluntary physical activity

We showed that K14Q-MOTS-c has a reduced effect on insulin sensitivity and weight gain compared to WT-MOTS-c in mice and speculate that the m.1382A>C polymorphism increases the prevalence of T2D in sedentary men because the C allele carriers produce a less effective K14Q-MOTS-c. However, an alternative explanation for the increased T2D frequency is that the m.1382A>C polymorphism can impact voluntary physical activity, which in turn affects T2D prevalence (although the frequency of the C allele was not statistically different between the tertile of physical activity in the J-MICC study as shown in Supplemental Table 4). We analyzed a voluntary wheel running activity for five days in male and female mice injected with WT-MOTS-c and K14Q-MOTS-c (7.5-mg/kg; BID; IP). Neither WT-MOTS-c or K14Q-MOTS-c had a significant effect on voluntary physical activity (Figure 6A-6F). Therefore, we ruled out the possibility that voluntary physical activity plays role in the m.1382A>C polymorphism (and MOTS-c variation) effect on T2D prevalence.

**Figure 6.**
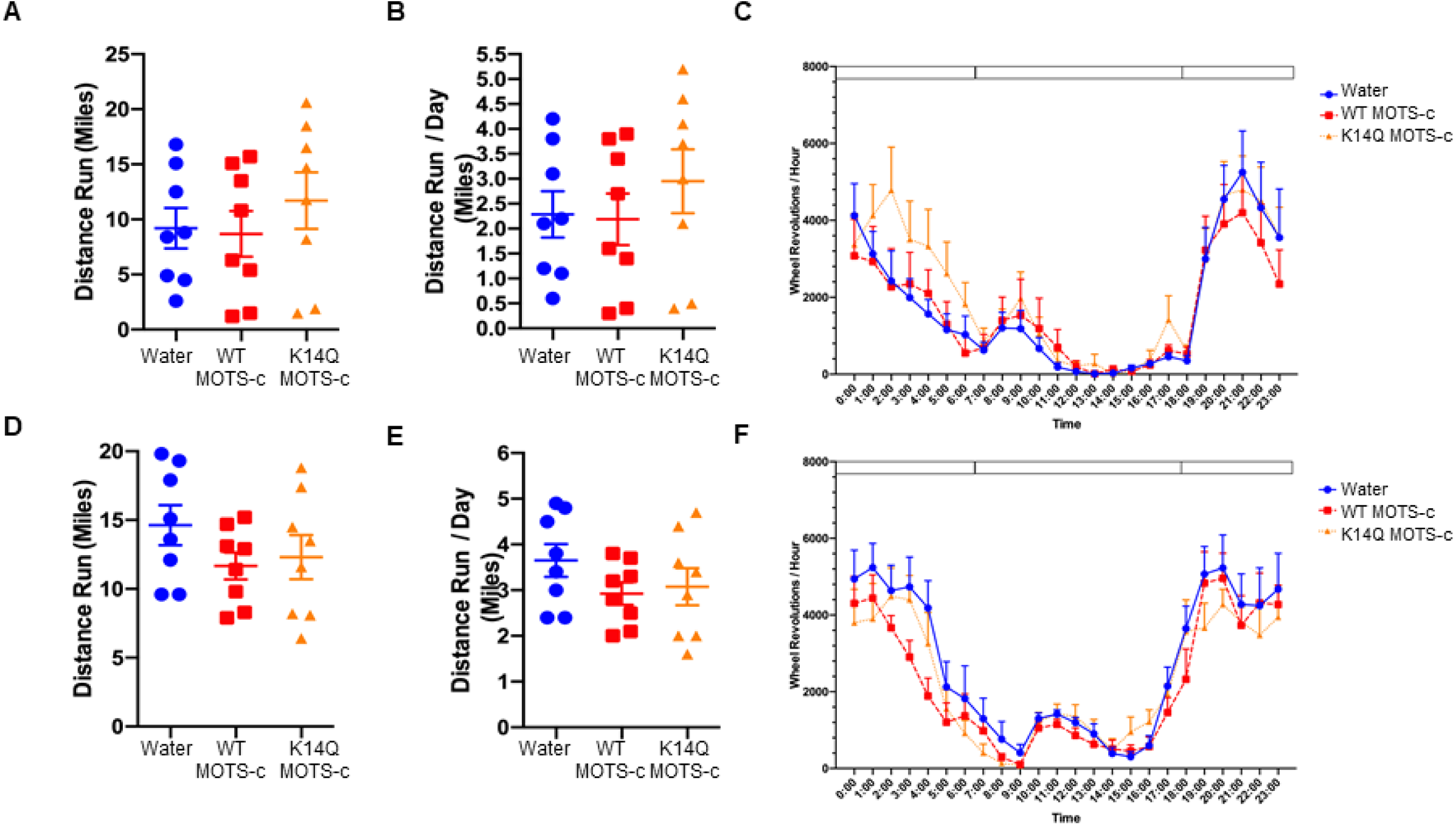
The effects of MOTS-c on voluntary wheel running activity. (A) total distance run, (B) total distance run per day, and (C) hourly running activity in males. (D) total distance run, (E) total distance run per day, and (F) hourly running activity in females.

## DISCUSSION

We found that C allele of the m.1382A>C polymorphism in the mtDNA results in the expression of an altered mitochondrial-derived peptide, K14Q-MOTS-c, and is associated with susceptibility to T2D, in sedentary Japanese men. The K14Q-MOTS-c peptide is less effective in promoting insulin sensitization and weight loss than wild type MOTS-c *in vitro* and *in vivo* studies. Thus, MOTS-c K14Q polymorphism might explain, at least in part, why East Asian populations such as Japanese, have and increased susceptibility to T2D despite their lower BMI compared with European populations. We also identified a sex difference in terms of the effects of the m.1382A>C polymorphism on T2D risk in humans and a related difference in the response to peptide administration in terms of glucose tolerance and weight gain in response to HFD feeding in mice. These observations represent a male-specific phenomenon. We noted that the levels of MVPA are related to the prevalence of T2D in C allele carrying men, representing a novel kinesio-genomic interaction. As the m.1382A>C polymorphism causes an amino acid replacement K14Q of MOTS-c peptide, we identified a reduced effect of K14Q MOTS-c in terms of its ability to induce metabolic improvements in high-fat diet-fed mice rather than modulating voluntary exercise.

In addition, our results suggest that higher MVPA rescues the potentially deleterious effects of the m.1382A>C polymorphism on T2D risk in men. As a result, exercise could be an especially important beneficial intervention for C allele carriers of this SNP. Developing an optimized exercise guidance for people with the m.1382A>C polymorphism, for example, could be clinically translatable.

It is attractive to hypothesize that MOTS-c is an exercise mimetic peptide that helps prevent T2D in sedentary men while in men who are frequent exercisers there is no further metabolic benefit of MOTS-c, or further protection from T2D by MOTS-c. On the other hand, the less active K14Q form may increase T2D risk, particularly in the susceptible sedentary group. We propose the concept that some variants of mtDNA evolved to best fit living conditions characterized by highly active lifestyles and limited caloric intake, where a thrifty gene such as K14Q-MOTS-c might present an advantage, but in the twenty-first century, it is a metabolic liability.

Our data shows that the effect of K14Q on T2D is male-specific. Hormonal differences between males and females could interact differentially with MOTS-c on metabolic function ^35^. Recently, MOTS-c was shown to prevents ovariectomy-induced metabolic dysfunction. MOTS-c did not change the metabolic phenotype in female mice with normal hormonal function (similar to our finding here), but MOTS-c improves the metabolic functions in ovariectomized female mice ^24, 25^. These results suggest the link of hormone and the effect of MOTS-c. Thus, it could be clinically relevant to identify what sex-specific factors permit females to overcome the potentially harmful effect of the m.1382A>C polymorphism on T2D seen in sedentary men. These may include sex-specific variation in mitochondrial mass or function or direct effects of sex-steroids on mitochondrial biology that favor females ^36^.

We believe these results are generalizable to the entire Japanese population and likely to all of East Asians. The prevalence of T2D in the present study was similar to the general Japanese population in their 50’s (men = 11.7%; women = 5.5%) ^37^, and the subject’s characteristics are representative of the mean BMI of the Japanese population in their late 50’s (men = 23.6 ± 1.3 kg/m^2^; women = 21.7 ± 1.1 kg/m^2^) ^38^.

The East Asian populations that may carry the MOTS-c K14Q SNP include around half a billion people in which the frequency of the m.1382A>C polymorphism varies from 5 to 8%. Sedentary behavior is alarmingly common in modern Asian cities ^39^. The reported rate of diabetes in these countries is well over 10%, and rising rapidly ^40^, suggesting that several million men with T2D carry the kinesio-genomic MOTS-c SNP. As MOTS-c analogues are currently in clinical development for the treatment of T2D complications, including Nonalcoholic steatohepatitis (NASH), the recognition of the m.1382A>C associated kinesio-genomic T2D risk may inform future clinical trials. On the other hand, this novel discovery suggests that additional ethnic-specific mtDNA polymorphisms, that might affect the structure or expression of MOTS-c or other MDPs, may be involved in metabolic disease risk.

In summary, we found that C allele of the m.1382A>C polymorphism increases the risk of T2D in Japanese men, especially in sedentary individuals. This polymorphism causes an amino acid replacement from Lys (K) to Gln (Q) at amino acid 14 in the MOTS-c peptide, which renders it a less potent insulin-sensitizer compared to WT-MOTS-c. Understanding the effects of this genetic polymorphism will provide a basis for developing physical activity strategies to maximize the benefits of exercise in T2D.

## Methods

### Epidemiological study

#### Japan Multi-Institutional Collaborative Cohort (J-MICC)

A cohort in which the relationship between genes and lifestyle was examined ^41^. This cross-sectional study consisted of 12,068 subjects in Saga City (men, 5,078; women, 6,990) who were between 40–69 years old. The Saga J-MICC Study was approved by the Ethics Committees of both Saga University Faculty of Medicine and Nagoya University Graduate School of Medicine. The study conformed to the principles outlined in the Declaration of Helsinki. Written informed consent was obtained from all subjects before their inclusion in the study. A variety of lifestyle measurements were recorded. The baseline survey was conducted from November 1, 2005 through December 22, 2007 ^42^. A self-administered questionnaire was used to collect data on smoking, dietary habits, current medication, disease history, and family history. Daily physical activity was objectively measured using an accelerometer (Life-Corder; Suzuken, Nagoya, Japan) as described in a previous study ^43^. Height and weight were measured to the nearest 0.1 cm and 0.1 kg, respectively. Body mass index (BMI) was calculated as the weight in kilograms divided by the square of the height in meters (kg/m^2^). Waist circumference was measured to the nearest 0.1 cm at the midpoint between the lower costal margin and the iliac crest using a calibrated measuring tape. The HbA_1c_ (%) level was measured in laboratories. The HbA_1c_ (%) value was converted from the Japan Diabetes Society (JDS) to the National Glyco-hemoglobin Standardization Program (NGSP) by using the following equation published by the JDS: NGSP (%) = 1.02 × JDS (%) + 0.25%^44^. T2D of the subjects was defined as the presence of either questionnaire, medication, or over 6.5% of HbA_1c_ level. Mitochondrial genetic variants were captured as described previously ^11, 45^. Briefly, mitochondrial polymorphisms were determined with sequence-specific oligonucleotide probes (G&G Science, Fukushima, Japan) by use of suspension array technology (Luminex 100).

#### Multiethnic Cohort (MEC)

MEC is a population-based prospective cohort study including approximately 215,000 men and women from Hawaii and California that is part of the Population Architecture using Genomics and Epidemiology (PAGE) study. All participants were 45-75 years of age at baseline, and primarily of 5 ancestries: Japanese Americans, African Americans, European Americans, Hispanic/Latinos, and Native Hawaiians. All cohort members completed baseline and follow-up questionnaires. For the current study, MEC contributed data from Japanese American with T2D and case-control subjects (1,799 cases and 1,588 controls), both men and women with complete covariate data. Single variant association testing was completed among 3,387 MEC Japanese Americans with complete genotype data for the m.1382A>C variant (rs111033358). The variant was coded as haploid with individuals carrying 0/0.5/1 copies, across the sample set. Rather than removing the one individual identified as a heterozygous carrier, the variant was coded as “0.5” for that person for inclusion. A logistic regression model adjusted for age and BMI was used to identify stratified gender specific associations. All analyses were performed in SAS and adjusted beta values, odds ratios, 95% confidence intervals and p-values were reported.

#### Tohoku Medical Megabank Project (TMM)

Over 80,000 apparently healthy adults living in the Pacific coast of the Tohoku region in Japan were recruited from May 2013 to March 2016 for the TMM. The detail of the TMM was described elsewhere ^46^. We obtained approval from the relevant ethics committees. All participants gave written informed consent at the time of study enrolment. A total of 9,559 participants were genotyped using the HumanOmniExpressExome BeadChip Array (Illumina Inc., San Diego, CA, USA). 3,552 participants were genotyped by whole genome sequencing using HiSeq 2500 system, as described elsewhere ^47, 48^. The subjects answered their history of disorder, including that of T2D, in our research questionnaire. Association between mtDNA 1382A>C and T2D was tested by linear regression analysis using R version 3.5.1.

### Meta-analysis

In order to pool the results from the three cohorts, we performed a meta-analysis to compare the prevalence of T2D between m.1382A and C allele. Meta-analysis was conducted using the R package “metafor” ^49^. The DerSimonian and Laird random-effects model ^50^ were used and analysis was performed with odds ratio adjusted by age and BMI. The between population heterogeneity was assessed by using the I^2^ statistic ^51^, that a higher I^2^ values mean higher heterogeneity. Significance of the pooled effect was determined by the Z test, and 95% CIs were calculated.

### Molecular modeling

3D molecular modeling of MOTS-c 14K (MRWQEMGYIFYPR<u>K</u>LR) and 14Q (MRWQEMGYIFYPR<u>Q</u>LR) were carried out using the program PEP-FOLD3 (web server, http://bioserv.rpbs.univ-paris-diderot.fr/services/PEP-FOLD3) ^52^ and I-TASSER (web server, https://zhanglab.ccmb.med.umich.edu/I-TASSER/)^53^. PEP-FOLD3 predicts the folding characteristics of peptides comprising of 5 to 50 amino-acids. I-TASSER predicts the folding characteristics of peptides or proteins comprised of 10 to 1500 amino-acids. To predict 12S rRNA structure by m.1382 A>C polymorphism, we used RNAstructure (https://rna.urmc.rochester.edu/RNAstructureWeb/Servers/Predict1/Predict1.html). PyMOL was used to analyze the electro physical differences between m.1382 A and C.

### Cell culture

C2C12 mouse myoblasts and L6 rat myoblasts were purchased from ATCC (Cat#. CRL-1772 and CRL-1458; Manassas, VA, USA) and cultured in DMEM + 20% FBS and MEM +10% FBS, respectively (FBS; ThermoFisher Scientific, Waltham, MA, USA) at 37° C in 5% CO_2_. To differentiate C2C12 and L6 myoblasts to myotubes, media was replaced with DMEM + 2% horse serum every 24 h for 6-7 days until cells were fully differentiated. 3T3-L1 mouse embryonic fibroblasts were purchased from ATCC (Cat#. CL-173) and cultured in DMEM + 10% Bovine Calf Serum (pre-adipocyte expansion medium) at 37° C in 5% CO_2_. To differentiate 3T3-L1 pre-adipocytes, pre-adipocyte expansion medium was replaced with differentiation medium (DMEM,10% FBS, 1μM Dexamethasone, 0.5mM IBMX, 1μg/mL insulin) for 48 h, then replaced with adipocyte maintenance medium (DMEM 10% FBS, 1µg/mL insulin) every 2 days for 7 days. HEK293 cells were cultured in high glucose Dulbecco’s modified Eagle’s medium (DMEM; Sigma) supplemented with 10% fetal bovine serum at 37°C in 5% CO2. For transient transfection of the wild type (WT) and K14Q MOTS-c in pcDNA3.1(+), lipofectamine 3000 was used according to the manufacturer’s instruction (ThermoFisher Scientific).

### Insulin signaling

In order to examine insulin signaling, fully differentiated C2C12 media was replaced with low glucose DMEM + 0.1% fatty acid free (FAF)- BSA for 2hr at 37° C. Then, the media was replaced with PBS + 0.1% FAF-BSA for 30-min and 10-nM bovine insulin was added to the cells with or without peptides for 15 min at 37° C. After the incubation, cells were washed with PBS, and protein was extracted. Total and phosphor-AKT levels were measured by MSD (MESO SCALE DISCOVERY, Rockville, MD, USA).

### Nile Red staining

Seven days after 3T3-L1 cells were differentiated, lipid droplets were quantified using the Lipid Droplets Fluorescence Assay Kit (Cat#. 500001; Cayman chemical; Ann Arbor, MI, USA) according to the manufacturer’s instruction. Briefly, cells were plated and differentiated in a black clear-bottom 96-well plate. Cells were fixed for 10min at RT, and stained with Nile Red staining solution for 15min at RT. After washing the staining solution, the fluorescence intensity of lipid droplets read with filter sets designed to detect FITC (ex/em 485/535 nm).

### Measurements of glucose levels in culture media

Extracellular glucose from cell culture medium was measured using glucose assay kits per manufacturer’s instructions (Eton Biosciences, USA). Briefly, standard glucose solution and cell culture medium were incubated with glucose enzyme mix for 30min at 37° C. During the incubation, glucose is oxidized by enzyme reactions to yield a colored product, which can be measured at 570nm. After 30min, the absorbance was measured on a plate spectrophotometer (Molecular Designs, Sunnyvale, CA) at 570 nm. We calculated the value of glucose in the medium using the equation obtained from the linear regression of the standard curve.

### Animals

Male and female CD-1 mice at 14 weeks (n=8-12) and 12 weeks (n=8-11) of age, respectively, were obtained from Jackson Laboratory (Bar Harbor, ME USA) and treated with K14Q MOTS-c and WT MOTS-c peptides. Mice were singly housed under standard 12-hr light-dark cycle with access to water and rodent food a high-fat diet (HFD, 60% by calories; Research Diets, MO). Mice were randomly assigned to one of three experimental groups: 1) a control group receiving BID (twice a day) intraperitoneal (IP) injection of vehicle (sterilized water); 2) a WT MOTS-c-treated group receiving BID IP injection of 7.5mg per kg body weight; 3) a K14Q MOTS-c-treated group receiving BID IP injection of 7.5mg per kg body weight. Mice were anesthetized with isoflurane and sacrificed after two weeks of treatment. Tissues were collected from the mice, flash-frozen in liquid nitrogen and stored at −80°C. For glucose tolerance test (GTT), 12-week old male and female CD1 mice were treated with WT MOTS-c (0.5-mg/kg/day; IP), K14Q MOTS-c (0.5-mg/kg/day; IP), or sterile pure water (vehicle), daily for 21 days and blood glucose was measured using a glucometer (Freestyle, Abbott Laboratories; Abbott Park, IL, USA). GTT consisted of a D-glucose injection (1g/kg; IP) and blood was sampled from the tail at 0, 15, 30, 60, 90, and 120 minutes post-glucose injection. In-cage Voluntary Wheel Running Studies: Voluntary wheel running studies were performed on ten-week-old male and female CD-1 mice purchased from Charles River Laboratories. Animals were individually housed for the duration of the experimental protocol. Animals of both sexes were weighed and divided into three groups a priori to ensure equal body weights between groups. At twelve weeks of age, all animals were provided ad libitum access to a high fat diet (HFD, 60% kcal Fat; D12492, Research Diets). Two weeks after HFD initiation, all animals began receiving intraperitoneal injections of vehicle (sterilized water), MOTS-c, or K14Q MOTS-c (7.5 mg/kg, BID). Four days after injection initiation, in cage running wheels were introduced to all animals. Five days after running wheel introduction, the animals received a final vehicle or treatment injection at the same time the running wheels were locked. After the wheels were locked, all mice were fasted for 6 hours prior to euthanasia and tissue removal (i.e. whole blood, quadriceps muscle, gonadal white adipose tissue, and liver). All experiments with mice were performed in accordance with the appropriate guidelines and regulations and approved by the Juntendo University, University of Southern California, and University of California, Los Angeles Institutional Animal Care and Use Committee.

### Statistical Analysis

Statistical analyses were performed using PASW Statistics 18 (series of SPSS Statistics) for Windows. Differences in physical characteristics by sex were determined using unpaired t-tests and Pearson’s χ^2^ tests. The relation of the m.1382A>C genotype distribution and prevalence of T2D was evaluated using the Pearson’s χ^2^ test. Logistic regression was also used to predict the prevalence of T2D. All logistic regression models included the following covariates: age, BMI, smoking, and moderate-to-vigorous intensity physical activity (MVPA). The statistical significance for linearity between the prevalence of T2D and MVPA was evaluated using Cochran-Armitage trend tests. In animal and cell culture studies, group comparison was performed by t-test or one-way analysis of variance (ANOVA). The Bonferonni post hoc test was performed when the ANOVA indicated a significant difference. Delta body weight, fat mass, lean mass and glucose tolerance test were analyzed with two-way (group × time) ANOVA with replications. Values are expressed as mean ± SD; *p* < 0.05 is considered to be statistically significant. Cell and mouse data are presented as mean ± S.E.M. Significant differences were determined by Student’s t-tests or one-way ANOVA and Turkey’s post hoc test by use of GraphPad Prism 5 software. P values of *<0.05, **<0.01, or ***<0.001 were considered statistically significant.

## Acknowledgments

This study was supported in part by Japan Society for Promotion of Science (JSPS) KAKENHI Grants (17015018, 221S0001, 16H06277, and 17H01554 to K.T.; 16K09058 to M.H.; 16K13052 to N.F.; and 18K17943 to H.Z.) and by the MEXT-Supported Program for the Strategic Research Foundation at Private Universities (to Juntendo University). H.K. was a recipient of a Grant-in-Aid for JSPS fellows (17J10817). This work was also supported by a Glenn/AFAR Postdoctoral Fellowship Program for Translational Research on Aging to S.J.K and by grant P01AG034906 and U54CA233465 to P.C.

## Author contributions

H.Z. and S.J.K. are joint first author to write an article. Y.N., Y.H., M.H., K.T., Y.S., A.S., K.K., K.K., Y.M.P. and V.W.S. organized human study cohort. N.F., Y.N., and E.M-M. extracted and analyzed DNA. H.Z., S.J.K. and N.F. performed statistical analysis and wrote an article. H.Z., H.K., E.M-M., H.N., and N.F. performed mice experiments. S.J.K. perform a majority of experiments related to cells and mice. J. W. is a director of the aging biomarker and measured P-AKT/T-AKP levels using MSD. K.Y., B.M., J.X., and H.H.M. contributed mice experiments. R.V. contributed 3T3-L1 experiments. C.L. contributed MOTS-c plasmid experiments. Y.M.P. and V.W.S. analyzed the P.C. and N.F. supervised the project and S.J.K., H.Z., N.F. and P.C. finalized this article.

## Supplemental information

Supplemental Figure 1. (A-B) The models were obtained via the program PEP-FOLD3 (model 1). The left two structures show the model of MOTS-c 14K peptide. The right two structures show the model of MOTS-c 14Q peptide. The lower view is generated by rotating peptide structure 90°around the y-axis from upper view. (C-D) The 3D structure of MOTS-c 14K and 14Q models obtained via the program I-TASSER (model 2). The figure is generated by Molmil viewer (https://pdbj.org/molmil/).

Supplemental Table 1. Characteristics of J-MICC subjects by the m.1382A>C polymorphism.

Supplemental Table 2. Characteristics of TMM subjects by the m.1382A>C polymorphism.

Supplemental Table 3. Association of m.1382C with T2D among MEC Japanese Americans-haploid analysis.

Supplemental Table 4. Characteristics of J-MICC subjects by m.1382A>C polymorphism divided with physical activity.

Supplemental Table 5. Characteristics of J-MICC subjects by the m.4883C>T polymorphism (haplogroup D).

Supplemental Table 6. Characteristics of J-MICC subjects by the m.5178C>A polymorphism (haplogroup D).

Supplemental Table 7. Characteristics of J-MICC subjects by the m.3010C>A polymorphism (haplogroup D4).

Supplemental Table 8. Characteristics of J-MICC subjects by the haplogroup D4b (m.1382C or m.15440C).

Supplemental Table 9. Factors associated with the risk of type 2 diabetes in the J-MICC subjects divided with moderate-vigorous physical activity.

