## Supplemental Tables and Figures for "A Pro-Diabetogenic mtDNA Polymorphism in the Mitochondrial-Derived Peptide, MOTS-c"

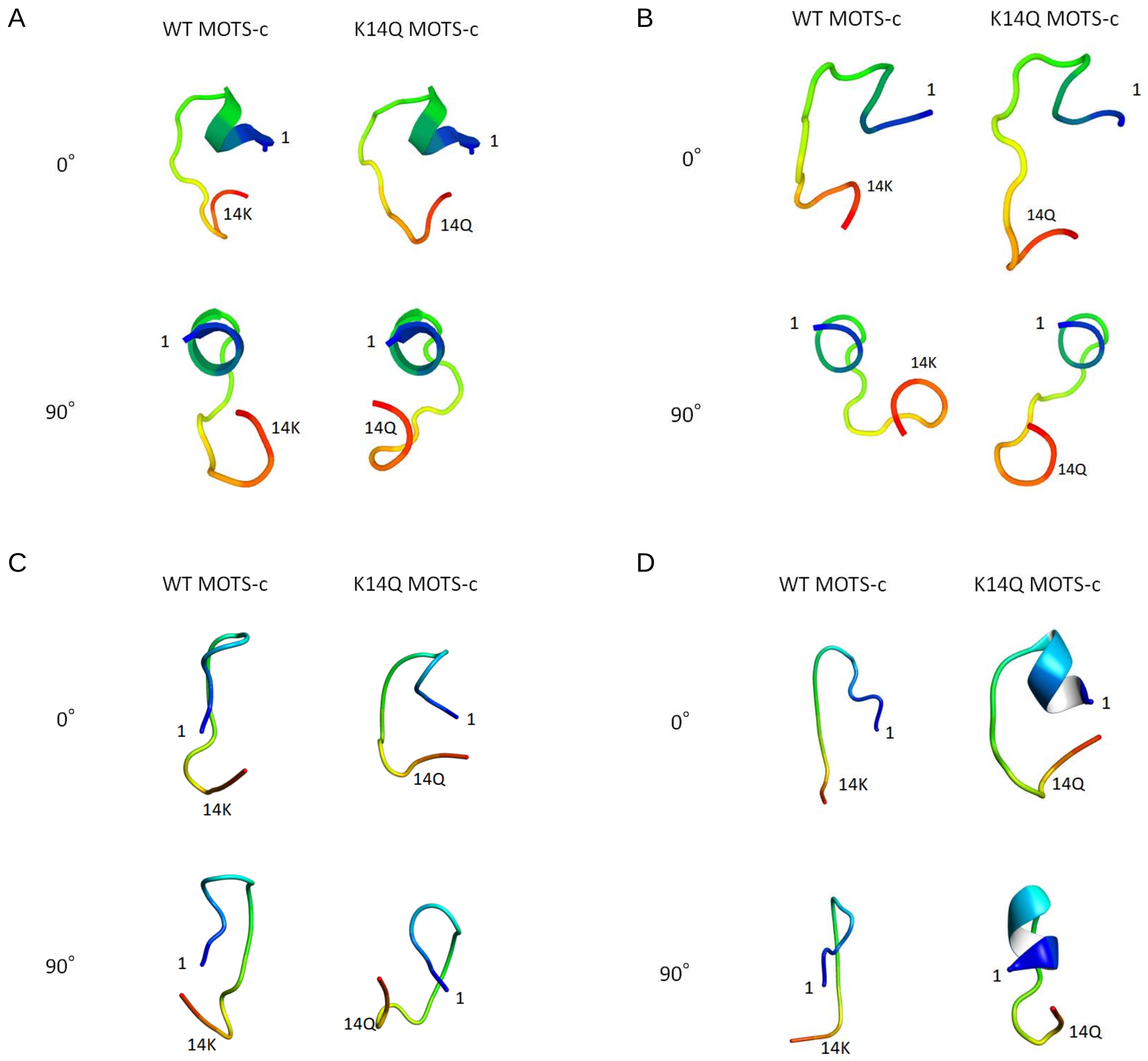

**Supplemental Figure 1.** (A-B) The models were obtained via the program PEP-FOLD3 (model 1). The left two structures show the model of MOTS-c 14K peptide. The right two structures show the model of MOTS-c 14Q peptide. The lower view is generated by rotating peptide structure 90°around the y-axis from upper view. (C-D) The 3D structure of MOTS-c_14K and 14Q models obtained via the program I-TASSER (model 2). The figure is generated by Molmil viewer (https://pdbj.org/molmil/).

**Supplemental Table 1.** Characteristics of J-MICC subjects by the m.1382A>C polymorphism.

| | A | C |
|---|---|---|
| Male | | |
| n | 4,603 | 360 |
| Age (years old) | 56.6 ± 8.2 | 55.8 ± 8.6 |
| Height (cm) | 166.9 ± 6.0 | 167.3 ± 5.6 |
| BMI (kg/m$^2$) | 24.2 ± 3.0 | 24.3 ± 3.3 |
| Waist circumference (cm) | 86.2 ± 7.9 | 86 ± 8.7 |
| $HbA_{1c}$ (%) | 5.3 ± 0.8 | 5.3 ± 0.9 |
| Moderate to vigorous physical activity (min/day) | 21.6 ± 17.5 | 22.2 ± 21.3 |
| Smokers (%) | 35.7 | 38.6 |
| Prevalence of type 2 diabetes (%) | 10.8 | 13.1 |
| Female | | |
| n | 6,356 | 534 |
| Age (years old) | 55.6 ± 8.2 | 55.8 ± 8.3 |
| Height (cm) | 154.5 ± 5.4 | 154.5 ± 5.7 |
| BMI (kg/m$^2$) | 22.8 ± 3.2 | 22.9 ± 3.4 |
| Waist circumference (cm) | 81.2 ± 9.4 | 81.4 ± 9.8 |
| $HbA_{1c}$ (%) | 5.2 ± 0.6 | 5.2 ± 0.6 |
| Moderate to vigorous physical activity (min/day) | 18.8 ± 13.5 | 17.7 ± 12.7 |
| Smokers (%) | 8.4 | 9.4 |
| Prevalence of type 2 diabetes (%) | 4.7 | 3.7 |

BMI, body mass index. Mean ± SD.

**Supplemental Table 2.** Characteristics of TMM subjects by the m.1382A>C polymorphism.

| | A | C |
|---|---|---|
| Male | | |
| n | 4,199 | 272 |
| Age (years old) | 61.4 ± 11.5 | 60.8 ± 12.6 |
| Height (cm) | 162.9 ± 11.1 | 162.0 ± 12.6 |
| BMI ($kg/m^2$) | 23.4 ± 4.4 | 23.4 ± 5.0 |
| Waist circumference (cm) | 83.1 ± 11.5 | 83.1 ± 13.0 |
| $HbA_{1c}$ (%) | 5.6 ± 0.7 | 5.6 ± 0.8 |
| Smokers (%) | 25.6 | 24.3 |
| Prevalence of type 2 diabetes (%) | 12.4 | 15.4 |
| Female | | |
| n | 7,338 | 479 |
| Age (years old) | 58.0 ± 12.7 | 57.8 ± 13.1 |
| Height (cm) | 151.4 ± 8.7 | 150.5 ± 9.6 |
| BMI ($kg/m^2$) | 22.4 ± 4.6 | 22.0 ± 4.9* |
| Waist circumference (cm) | 79.5 ± 11.7 | 78.3 ± 12.6 |
| $HbA_{1c}$ (%) | 5.5 ± 0.5 | 5.5 ± 0.7 |
| Smokers (%) | 5.7 | 6.1 |
| Prevalence of type 2 diabetes (%) | 5.4 | 6.3 |

BMI, body mass index. *$p < 0.05$, vs A allele. Mean ± SD.

**Supplemental Table 3.** Association of m.1382C with T2D among MEC Japanese Americans-haploid analysis.

| | N (Cases/Controls) | m.1382C Freq | Beta * | OR* | 95% CI* | *p* value* |
|---|---|---|---|---|---|---|
| Males | 1,810 (1,016/794) | 0.076 | 0.3894 | 1.48 | 1.01-2.15 | 0.0424 |
| Females | 1,577 ( 783/794) | 0.068 | -0.188 | 0.83 | 0.53-1.28 | 0.4014 |

* The results above are adjusted for age and BMI.

**Supplemental Table 4**. Characteristics of J-MICC subjects by m.1382A>C polymorphism divided with physical activity.

| MVPA tirtile | T1 | | | T2 | | | T3 | | |
|---|---|---|---|---|---|---|---|---|---|
| m.1382A>C | A | C | *p* | A | C | *p* | A | C | *p* |
| Male | | | | | | | | | |
| (n) | 1,529 | 124 | | 1,536 | 119 | | 1,538 | 117 | |
| Age (years old) | 58.4 ± 0.2 | 57.8 ± 0.8 | 0.443 | 55.3 ± 0.2 | 53.9 ± 0.7 | 0.070 | 56.2 ± 0.2 | 55.6 ± 0.8 | 0.498 |
| Height (cm) | 167.2 ± 0.2 | 167.1 ± 0.6 | 0.909 | 167.4 ± 0.2 | 167.4 ± 0.5 | 0.955 | 166.2 ± 0.1 | 167.4 ± 0.5 | 0.034 |
| BMI (kg/m$^2$) | 24.3 ± 0.1 | 24.5 ± 0.3 | 0.605 | 24.2 ± 0.1 | 24.3 ± 0.3 | 0.726 | 24.1 ± 0.1 | 23.9 ± 0.3 | 0.461 |
| Waist circumference(cm) | 87.4 ± 0.2 | 87.2 ± 0.9 | 0.833 | 86.1 ± 0.2 | 85.7 ± 0.8 | 0.554 | 85.2 ± 0.2 | 85.0 ± 0.7 | 0.750 |
| HbA1c (%) | 5.3 ± 0.8 | 5.4 ± 0.8 | 0.222 | 5.3 ± 0.8 | 5.3 ± 0.7 | 0.951 | 5.3 ± 0.8 | 5.3 ± 1.1 | 0.456 |
| Smokers (%) | 43.1% | 43.5% | 0.497 | 35.3% | 41.2% | 0.117 | 28.7% | 30.8% | 0.349 |
| Prevalence of type 2 diabetes (%) | 11.2% | 18.5% | 0.014 | 10.0% | 12.6% | 0.218 | 11.4% | 7.7% | 0.141 |
| Female | | | | | | | | | |
| (n) | 2,105 | 190 | | 2,115 | 183 | | 2,136 | 161 | |
| Age (years old) | 57.4 ± 0.2 | 57.2 ± 0.6 | 0.823 | 54.4 ± 0.2 | 55.2 ± 0.6 | 0.235 | 55.0 ± 0.2 | 55.0 ± 0.6 | 0.995 |
| Height (cm) | 154.6 ± 0.1 | 154.0 ± 0.4 | 0.153 | 155.0 ± 0.1 | 155.1 ± 0.4 | 0.774 | 154.1 ± 0.1 | 154.3 ± 0.4 | 0.589 |
| BMI (kg/m$^2$) | 23.1 ± 0.1 | 23.5 ± 0.3 | 0.102 | 22.5 ± 0.1 | 22.4 ± 0.2 | 0.617 | 22.6 ± 0.1 | 22.7 ± 0.2 | 0.978 |
| Waist circumference(cm) | 82.9 ± 0.2 | 83.5 ± 0.8 | 0.388 | 80.6 ± 0.2 | 80.2 ± 0.7 | 0.559 | 80.2 ± 0.2 | 80.3 ± 0.7 | 0.910 |
| HbA1c (%) | 5.2 ± 0.7 | 5.2 ± 0.5 | 0.143 | 5.2 ± 0.6 | 5.2 ± 0.6 | 0.441 | 5.2 ± 0.6 | 5.2 ± 0.7 | 0.781 |
| Smokers (%) | 9.8% | 13.2% | 0.091 | 8.2% | 9.3% | 0.349 | 6.9% | 5.0% | 0.225 |
| Prevalence of type 2 diabetes (%) | 5.1% | 3.7% | 0.258 | 3.7% | 4.4% | 0.377 | 5.3% | 3.1% | 0.151 |

MVPA, moderate vigorous physical activity. BMI, body mass index. Mean ± SD.

**Supplemental Table 5.** Characteristics of J-MICC subjects by the m.4883C>T polymorphism (haplogroup D).

| | C | T |
|---|---|---|
| Male | | |
| n | 3,108 | 1,855 |
| Age (years old) | 56.6 ± 8.2 | 56.4 ± 8.1 |
| Height (cm) | 166.9 ± 6.0 | 167.1 ± 6.0 |
| BMI ($kg/m^2$) | 24.2 ± 2.9 | 24.2 ± 3.1 |
| Waist circumference (cm) | 86.2 ± 7.9 | 86.2 ± 8.2 |
| $HbA_{1c}$ (%) | 5.3 ± 0.8 | 5.3 ± 0.9 |
| Moderate to vigorous physical activity (min/day) | 22.0 ± 18.0 | 21.1 ± 17.4 |
| Smokers (%) | 35.7 | 36.2 |
| Prevalence of type 2 diabetes (%) | 10.8 | 11.4 |
| Female | | |
| n | 4,262 | 2,628 |
| Age (years old) | 55.6 ± 8.2 | 55.6 ± 8.2 |
| Height (cm) | 154.6 ± 5.4 | 154.4 ± 5.4 |
| BMI ($kg/m^2$) | 22.8 ± 3.2 | 22.8 ± 3.2 |
| Waist circumference (cm) | 81.3 ± 9.4 | 81.2 ± 9.6 |
| $HbA_{1c}$ (%) | 5.2 ± 0.6 | 5.2 ± 0.6 |
| Moderate to vigorous physical activity (min/day) | 19.1 ± 13.7 | 18.1 ± 13.0* |
| Smokers (%) | 7.8 | 9.3* |
| Prevalence of type 2 diabetes (%) | 4.8 | 4.4 |

BMI, body mass index. *$p < 0.05$. Mean ± SD.

**Supplemental Table 6**. Characteristics of J-MICC subjects by the m.5178C>A polymorphism (haplogroup D).

| | C | A |
|---|---|---|
| Male | | |
| n | 3,112 | 1,851 |
| Age (years old) | 56.6 ± 8.2 | 56.4 ± 8.1 |
| Height (cm) | 166.9 ± 6.0 | 167.1 ± 6.0 |
| BMI ($kg/m^2$) | 24.2 ± 3.0 | 24.2 ± 3.0 |
| Waist circumference (cm) | 86.2 ± 7.9 | 86.2 ± 8.2 |
| $HbA_{1c}$ (%) | 5.3 ± 0.8 | 5.3 ± 0.9 |
| Moderate to vigorous physical activity (min/day) | 22.0 ± 18.0 | 21.1 ± 17.4 |
| Smokers (%) | 35.7 | 36.3 |
| Prevalence of type 2 diabetes (%) | 10.8 | 11.3 |
| Female | | |
| n | 4,265 | 2,625 |
| Age (years old) | 55.6 ± 8.2 | 55.6 ± 8.2 |
| Height (cm) | 154.6 ± 5.4 | 154.4 ± 5.4 |
| BMI ($kg/m^2$) | 22.8 ± 3.2 | 22.8 ± 3.2 |
| Waist circumference (cm) | 81.3 ± 9.4 | 81.2 ± 9.6 |
| $HbA_{1c}$ (%) | 5.2 ± 0.6 | 5.2 ± 0.6 |
| Moderate to vigorous physical activity (min/day) | 19.1 ± 13.7 | 18.1 ± 13.0* |
| Smokers (%) | 7.8 | 9.3* |
| Prevalence of type 2 diabetes (%) | 4.8 | 4.4 |

BMI, body mass index. *$p < 0.05$. Mean ± SD.

**Supplemental Table 7.** Characteristics of J-MICC subjects by the m.3010C>A polymorphism (haplogroup D4).

| | G | A |
|---|---|---|
| Male | | |
| n | 3,287 | 1,676 |
| Age (years old) | 56.6 ± 8.2 | 56.5 ± 8.1 |
| Height (cm) | 166.9 ± 6.1 | 167.1 ± 5.9 |
| BMI ($kg/m^2$) | 24.2 ± 2.9 | 24.2 ± 3.1 |
| Waist circumference (cm) | 86.2 ± 7.9 | 86.3 ± 8.2 |
| $HbA_{1c}$ (%) | 5.3 ± 0.8 | 5.3 ± 0.9 |
| Moderate to vigorous physical activity (min/day) | 22.0 ± 17.9 | 21.0 ± 17.4 |
| Smokers (%) | 35.8 | 36.0 |
| Prevalence of type 2 diabetes (%) | 10.9 | 11.2 |
| Female | | |
| n | 4,504 | 2,386 |
| Age (years old) | 55.6 ± 8.2 | 55.6 ± 8.3 |
| Height (cm) | 154.6 ± 5.4 | 154.4 ± 5.4 |
| BMI ($kg/m^2$) | 22.8 ± 3.2 | 22.8 ± 3.2 |
| Waist circumference (cm) | 81.2 ± 9.4 | 81.2 ± 9.6 |
| $HbA_{1c}$ (%) | 5.2 ± 0.6 | 5.2 ± 0.6 |
| Moderate to vigorous physical activity (min/day) | 19.0 ± 13.7 | 18.1 ± 13.1* |
| Smokers (%) | 7.8 | 9.5* |
| Prevalence of type 2 diabetes (%) | 4.8 | 4.3 |

BMI, body mass index. *$p < 0.05$. Mean ± SD.

**Supplemental Table 8.** Characteristics of J-MICC subjects by the haplogroup D4b (m.1382C or m.15440C).

| | Others | D4b |
|---|---|---|
| Male | | |
| n | 4,514 | 449 |
| Age (years old) | 56.6± 8.2 | 56.0± 8.5 |
| Height (cm) | 166.9± 6.0 | 167.2± 5.7 |
| BMI (kg/m$^2$) | 24.2± 3.0 | 24.3± 3.2 |
| Waist circumference (cm) | 86.2± 7.9 | 86.1± 8.6 |
| $HbA_{1c}$ (%) | 5.3± 0.8 | 5.3± 0.9 |
| Moderate to vigorous physical activity (min/day) | 21.7 ± 17.5 | 21.7 ± 20.4 |
| Smokers (%) | 35.7 | 37.6 |
| Prevalence of type 2 diabetes (%) | 10.9 | 12.5 |
| Female | | |
| n | 6,238 | 652 |
| Age (years old) | 55.6± 8.2 | 55.5± 8.2 |
| Height (cm) | 154.5± 5.4 | 154.5± 5.6 |
| BMI (kg/m$^2$) | 22.8± 3.2 | 22.9± 3.4 |
| Waist circumference (cm) | 81.2± 9.4 | 81.1± 9.8 |
| $HbA_{1c}$ (%) | 5.2± 0.6 | 5.2± 0.6 |
| Moderate to vigorous physical activity (min/day) | 18.8 ± 13.5 | 17.8 ± 12.7 |
| Smokers (%) | 8.2 | 9.8 |
| Prevalence of type 2 diabetes (%) | 4.7 | 3.5 |

BMI, body mass index. Mean ± SD.

**Supplemental Table 9.** Factors associated with the risk of type 2 diabetes in the J-MICC subjects divided with MVPA.

| MVPA tietile | | T1 | | | T2 | | | T3 | |
|---|---|---|---|---|---|---|---|---|---|
| Male | | | | | | | | | |
| | Age | 1.040 | (1.019-1.062) | Age | 1.082 | (1.058-1.106) | Age | 1.050 | (1.029-1.071) |
| | BMI | 1.070 | (1.021-1.122) | BMI | 1.070 | (1.014-1.130) | BMI | 1.074 | (1.016-1.134) |
| | m.1382A>C | 1.837 | (1.131-2.983) | Constant | 0.000 | | Constant | 0.001 | |
| | Constant | 0.002 | | | | | | | |
| Female | | | | | | | | | |
| | Age | 1.068 | (1.038-1.099) | Age | 1.089 | (1.056-1.123) | Age | 1.088 | (1.059-1.118) |
| | BMI | 1.168 | (1.120-1.219) | BMI | 1.217 | (1.144-1.295) | BMI | 1.083 | (1.018-1.151) |
| | Constant | 0.000 | | Constant | 0.000 | | Constant | 0.000 | |

MVPA, moderate-vigorous physical activity. Values are odds ratio (95%CI). All logistic regression models included the following covariates: age, BMI, smoking (yes), and D4b haplogroup SNPs [m.4883C>T (haplogroup D), m.5178C>A (haplogroup D), m.3010G>A (haplogroup D4), m.15440T>C (haplogroup D4b1), and m.1382A>C (haplogroup D4b2)].
